# Amphibian larvae benefit from a warm environment under simultaneous threat from chytridiomycosis and ranavirosis

**DOI:** 10.1101/2022.09.20.508725

**Authors:** Dávid Herczeg, Dóra Holly, Andrea Kásler, Veronika Bókony, Tibor Papp, Hunor Takács-Vágó, János Ujszegi, Attila Hettyey

## Abstract

Rising temperatures can facilitate epizootic outbreaks, but disease outbreaks may be suppressed if temperatures increase beyond the optimum of the pathogens while still within the temperature range that allows for effective immune function in hosts. The two most devastating pathogens of wild amphibians, Batrachochytrium dendrobatidis (Bd) and ranaviruses (Rv), co-occur in large areas, yet little is known about the consequences of their co-infection and how these consequences depend on temperature. Here we tested how co-infection and elevated temperatures (28 and 30°C vs. 22°C) affected Bd and Rv prevalence, infection intensities, and resulting mortalities in larval agile frogs and common toads. We found multiple pieces of evidence that the presence of one pathogen influenced the prevalence and/or infection intensity of the other pathogen in both species, depending on temperature and initial Rv concentration. Generally, the 30°C treatment lowered the prevalence and infection intensity of both pathogens, and, in agile frogs, this was mirrored by higher survival. These results suggest that if temperatures naturally increase or are artificially elevated beyond what is ideal for both Bd and Rv, amphibians may be able to control infections and survive even the simultaneous presence of their most dangerous pathogenic enemies.

## Introduction

During recent decades amphibians have experienced dramatic declines [1] and have become one of the most threatened vertebrate taxa, pushing them into the forefront of conservation efforts [2]. The causes behind population declines and extinctions are complex [3], with multiple stressors acting in synergy, but emerging infectious diseases likely play a decisive role [4]. The chytrid fungus *Batrachochytrium dendrobatidis* (*Bd*) was linked directly to the extinction of dozens of amphibian species and caused the worst pandemic of wildlife ever recorded in history [1, 4]. At the same time, epidemics caused by Ranaviruses (family *Iridoviridae*, hereafter *Rv*) have also led to mass mortality events, resulting in local extinctions of amphibians [5-7].

Typically, several pathogens are present in natural populations [8], and these are likely to interactively determine consequences for hosts. Accordingly, co-infections are increasingly recognised as important drivers of disease dynamics [9, 10]. Co-infections can be disadvantageous, insignificant or even beneficial to hosts, and multiple levels of interactions can influence the outcomes [11]. Direct and indirect interactions among infectious agents within hosts include interference competition for physical space [12], exploitation competition for resources [13, 14], as well as indirect interactions mediated by cross-reaction immunity and the immunosuppression of the host [15, 16]. The outcome of co-infections is also shaped by the species-specific infectivity and virulence of pathogens [17], arrival order (i.e., ‘priority effect’ [18, 19]), species and body condition of hosts [20, 21], and abiotic factors [11].

The highly pathogenic and widespread *Bd* and *Rv* frequently co-infect amphibians [8, 22-24], and this can cause mass die-offs [25]. Naturally co-infected amphibians tend to experience higher *Bd* infection loads during simultaneous infection with *Rv* than individuals exclusively infected with *Bd* [22], while *Rv* infection intensity tends to be negatively associated with the probability of infection with *Bd*, but this relationship depends on frog taxa [24]. Nonetheless, experimental studies scrutinising the effects of co-infection with *Bd* and *Rv* on mortality, morbidity, pathogen prevalence and infection intensity are extremely scarce. The only such study to our knowledge [26] found non-additive effects of co-infection on pathogen loads but not on host growth, survival, and antibody response in post-metamorphic Cuban tree frog *Osteopilus septentrionalis*.

Co-infection with *Bd* and *Rv* occurs even though their thermal requirements are different. The optimum temperature range of *Bd* falls between 17°C and 25°C [27], and its critical thermal maximum lies around 28°C. Therefore, *Bd*-prevalence [28, 29] and mortality attributable to *Bd* infection [30] are usually highest during moderately warm months and in habitats where the temperature does not become exceedingly high [31, 32], while high temperatures promote host survival [30]. In contrast, the type species of the genus *Ranavirus, Frog Virus 3* (FV3), replicates successfully *in vitro* between 8 and 30°C, with a lower replication rate below 15°C and the highest rate at 30°C [33]. Indeed, deaths caused by *Rv* are more frequent during the warm summer months [34], so temperature also appears to be a crucial determinant of ranavirosis dynamics [35].

Temperature influences infection outcome not only through its effects on the growth of pathogens but also via its influence on amphibian hosts [11]. The positive temperature dependence of the immune system of amphibians [36, 37] may at least partly explain why individuals that occur in warmer areas can keep *Bd* infection at bay [38-40]. Accordingly, thermal treatments have proven helpful for clearing the *Bd* infection of amphibians in the laboratory [41-43]. However, a simple warmer-is-better rule does not apply universally to amphibian species facing chytridiomycosis, as the immune system of cold-adapted amphibians may be more effective against *Bd* at lower than at higher temperatures [44-46]. In the case of ranavirosis, elevating the temperature from 20 to 27°C increased *Rv* propagation, disease incidence, and mortality rate in the common frog *Rana temporaria* [35].

Similarly, an increase in temperature from 10 to 25°C resulted in higher mortality and *Rv* copy numbers in tadpoles of four amphibian species exposed to FV3 [47]. In contrast, *Rv* infection probability and mortality were lower at 22°C than at 14°C in two species of *Lithobates* frogs infected with three different FV3 strains [48]. While these studies delivered clear evidence for the importance of temperature in the case of both chytridiomycosis and ranavirosis, contradictions in patterns may be due to interspecific differences in the temperature dependence of amphibian immune functions or to some other factor remaining to be explored. Importantly, manipulative studies testing how elevated temperature affects disease progression in amphibians co-infected with these two pathogens are lacking entirely.

Hence, to clarify the effects of high temperatures on disease progression in amphibians during single and co-infections with *Bd* and *Rv*, we experimentally infected tadpoles of agile frogs (*Rana dalmatina*) and common toads (*Bufo bufo*) and subsequently exposed them to elevated temperatures for six days, and finally assessed infection patterns and survival. We thereby aimed to deliver information about the effects of high temperatures that can occur under natural conditions on the severity of consequences of *Bd* and *Rv* infection and, especially, on the outcomes of co-infection. We also wanted to provide information for the parameterisation of predictive models on the consequences of climate change on future dynamics of wildlife diseases. Finally, we intended to test the potential of localised heating, an *in situ* mitigation method relying on the thermal treatment of *Bd*-infected amphibians [49], by assessing whether elevated temperatures decrease *Bd* prevalence and intensity without resulting in elevated *Rv* infection loads and excessive mortality in the presence of *Rv*.

## Methods

### Experimental design and procedures

We applied a full-factorial design with three thermal treatments: 22, 28 and 30°C, combined with six infection treatments: uninfected control (‘control’); exposed to *Bd* (‘*Bd*’); exposed to *Rv* in a low concentration (‘*Rv*-low’); exposed to *Rv* in a high concentration (‘*Rv*-high’); co-exposed to *Bd* and *Rv* in a low concentration (‘*Bd* + *Rv*-low’); co-exposed to *Bd* and *Rv* in a high concentration (‘*Bd* + *Rv*-high’). We replicated each treatment combination 20 times (two individuals from each of ten families in each treatment combination) for a total of 360 animals per species. The experimental procedure started with a 19-days *Bd* treatment, followed by a 24-hours *Rv* treatment and, finally, a 6-days thermal treatment (for a schematic representation, see Fig. 1).

**Fig. 1.**
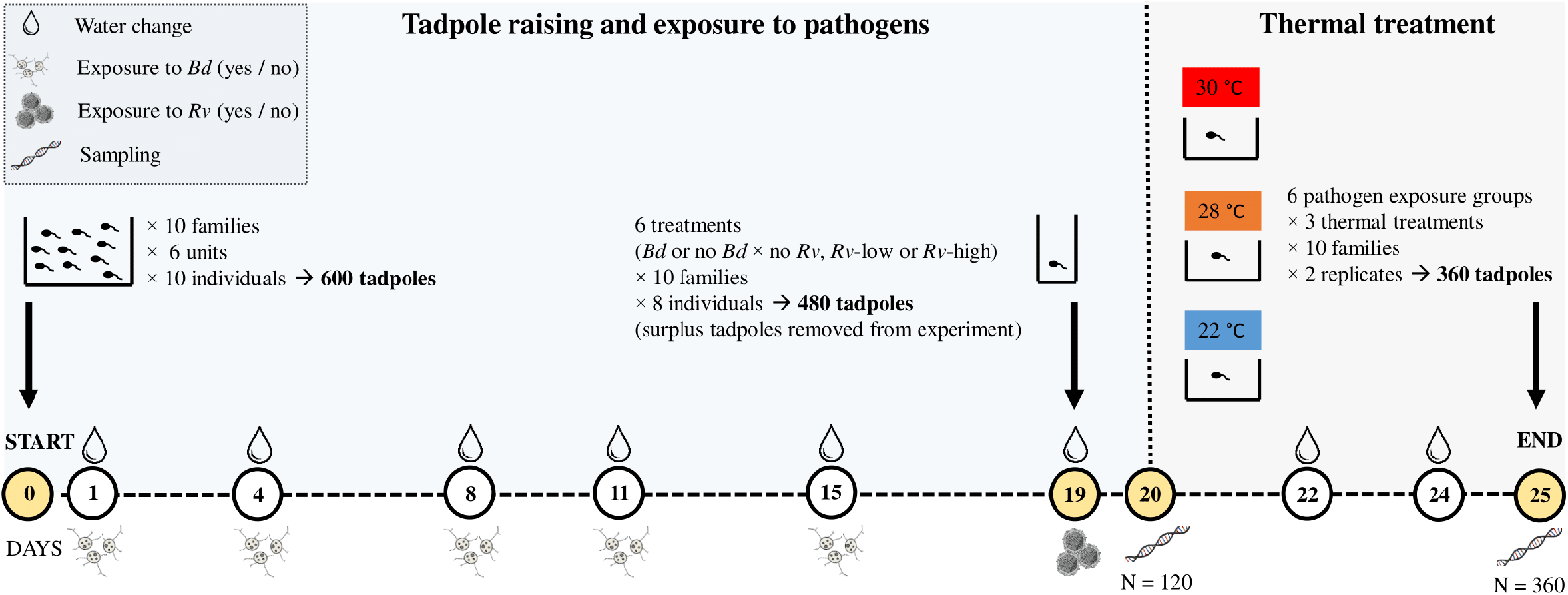
Schematic representation of the experimental procedure. *Bd* = *Batrachochytrium dendrobatidis*; *Rv* = *Ranavirus*

We collected 100 eggs from each of ten freshly laid clutches (families) from two natural populations of both species and transported them to the laboratory, where we reared embryos until hatching. Five days after hatching (development stage 25 according to Gosner [50]), we divided sibships into groups of 10 larvae and placed tadpoles into plastic rearing containers holding 10 litres of reconstituted soft water (RSW [51]; for details on animal collection and husbandry, see the Supplementary material). We randomly assigned rearing containers to *Bd* treatments, with one container representing each treatment by family combination. We performed the first infection with *Bd* on day 1 and, after that, renewed *Bd* concentrations following each water change until performing infections with *Rv*, which resulted in a total of five occasions of *Bd*-addition. On each occasion, we exposed tadpoles to approximately 2000 zoospores×ml^-1^ of *Bd* directly in their rearing containers, while the control tadpoles received sterile broth (for a detailed description of experimental infection with *Bd*, see the Supplementary material).

On day 19, we haphazardly selected eight tadpoles from each rearing container and randomly assigned them to *Rv* × temperature combinations. Then we exposed tadpoles to FV3 by applying one of two concentrations, i.e., *Rv*-low (6.12×10^3^ plaque forming unit (pfu)×ml^-1^) and *Rv*-high (6.25×10^5^ pfu×ml^-1^) during 24-hour exposures where tadpoles were challenged individually in plastic cups containing RSW and the corresponding concentration of *Rv*, while control tadpoles received sham extract (for more details on experimental infection with *Rv*, see the Supplementary material). Subsequently, to assess initial infection prevalence and intensity for both pathogens, we haphazardly selected 20 tadpoles from each of the six infection treatment groups (12 tadpoles from each family, 120 individuals in total) and preserved them in 96% ethanol.

On day 20, we started thermal treatments as described in Ujszegi et al. [52] (for details, including slight modifications, see the Supplementary material) and monitored tadpoles daily to record any mortality events. Water temperatures were kept at 22, 28, and 30 °C in the three thermal treatment groups, respectively (Table S1). Six days after the start of thermal treatments, we terminated the experiment by preserving all surviving tadpoles in 96% ethanol.

We extracted *Bd* DNA from dissected mouthparts and *Rv* DNA from liver tissue. We assessed infection prevalence and intensity using qPCR following standard amplification methodologies (see [53] for *Bd* and [54] for *Rv*). When the qPCR result was equivocal, we repeated reactions in duplicate. If we obtained an equivocal result again, we considered the sample positive [55] (for more details, see the Supplementary material).

### Statistical analyses

We analysed data on the two species separately. In the main text, we focus on infection status after the thermal treatment; see the Supplementary material for analyses before thermal treatment. Because of low *Bd* prevalence in agile frogs and low variance in infection intensities of both pathogens in common toads (see Table S2 and Table S3), we did not perform statistical modelling in these three cases. To analyse the effects of co-infection, thermal treatment, and their interaction on pathogen prevalence after temperature treatment, we used generalised linear mixed-effects models (GLMM) with binomial error, including only those treatment groups that had been exposed to the given pathogen. To analyse the effect of treatments and their interactions on *Rv* infection intensity after temperature treatment within *Rv*-positive agile frogs, we used a linear mixed-effects model (LMM). We analysed the survival of agile frog tadpoles during thermal treatment using a mixed-effects Cox’s proportional hazards model (COXME); mortality of common toads was negligible (1.1%; see Table S3), so we did not analyse it. In all models, we used family as a random factor. After stepwise model simplification, we performed pairwise comparisons by calculating linear contrasts and applying false discovery rate correction. For further details regarding the statistical analyses, see the Supplementary material.

## Results

### Agile frogs

Initial *Bd*-prevalence, as assessed before thermal treatments, was extremely low: two out of 60 *Bd*-exposed individuals carried the fungus. Initial *Bd* infection intensities were also very low (Table S2). In contrast, the initial prevalence of *Rv* in *Rv*-exposed tadpoles was between 73.7 (Table S2) and 100% and *Rv* infection intensities were low to moderate (Fig. S1; for details, see the Supplementary material).

By the end of the 6-days thermal treatments, *Bd* prevalence in *Bd*-exposed agile frog tadpoles remained low, with only three tadpoles testing positive, all in the *Bd* + *Rv*-low treatment (Table S3). Infection intensity of *Bd* was also very low after thermal treatments (<12.2 Genomic Equivalents (GE); Table S3).

The prevalence of *Rv* in *Rv*-exposed tadpoles after thermal treatments varied between 21.4 and 100% (Table S3; Fig. 2A) and was higher in the *Rv*-high treatment (GLMM; χ^2^_1_ = 14.49, *P* < 0.001) and in tadpoles not co-exposed to *Bd* (χ^2^_1_ = 13.34, *P* < 0.001). There was a tendency for thermal treatment to affect *Rv* prevalence (χ^2^_2_ = 4.62, *P* = 0.099), where the lowest infection probability was at 28 and the highest at 22°C (Table S4). The two-way interactions were non-significant (all *P* > 0.37).

**Fig. 2.**
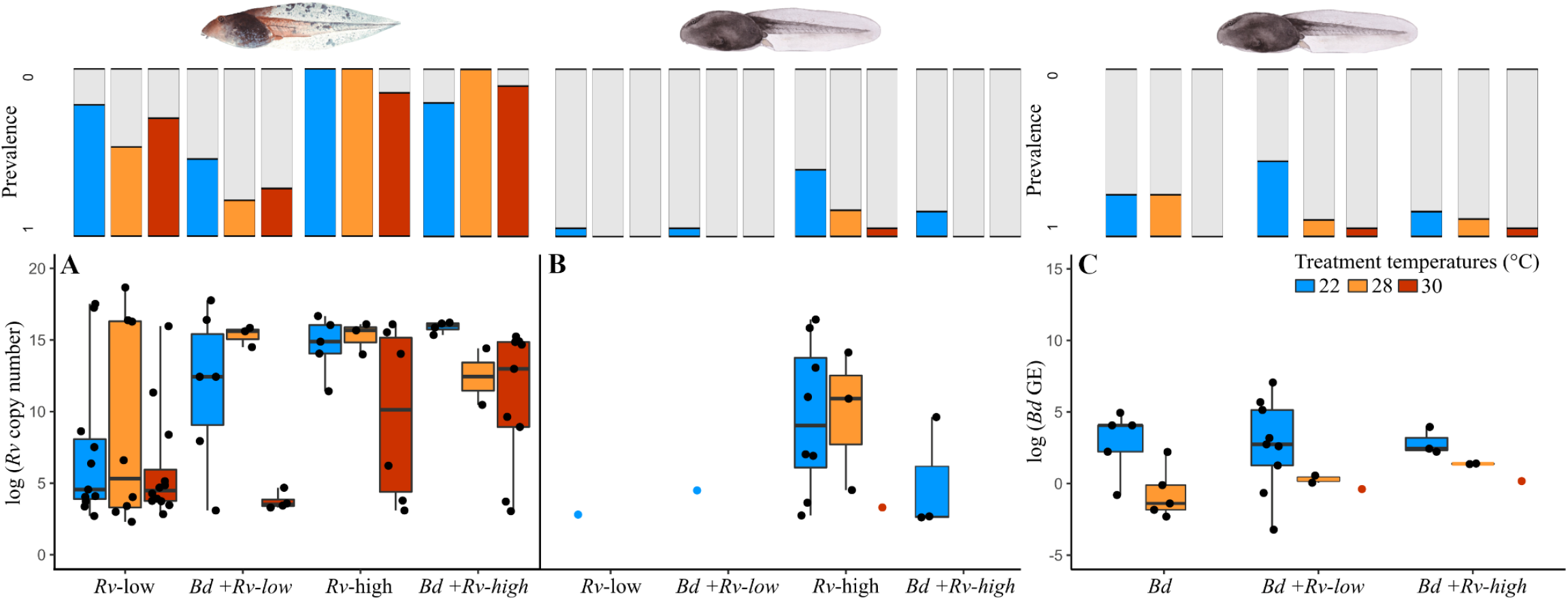
(A) *Ranavirus* (*Rv*) infection intensities in *Rv*-positive agile frog tadpoles with corresponding prevalences and (B and C) *Rv* and *Batrachochytrium dendrobatidis* (*Bd*) infection intensities in *Rv*- and *Bd*-positive common toads with corresponding prevalences of pathogens after the 6-days of thermal treatment, following exposure to infection treatments. The *Rv* and *Bd* infection intensity data (black dots) were natural log-transformed. Treatment groups with only one intensity data points represented as coloured dots. Horizontal lines represent medians, boxes represent interquartile ranges, and whiskers represent minimum-maximum ranges.

*Ranavirus* infection intensities were high and were positively affected by *Rv* concentration (LMM; χ^2^_1_ = 15.34, *P* < 0.001), whereas the main effects of thermal treatment (χ^2^_2_ = 4.34, *P* = 0.11) and previous exposure to *Bd* (χ^2^_1_ = 2.24, *P* = 0.14) were non-significant. The two-way interactions were all non-significant as well (all *P* > 0.17), but the three-way interaction between thermal treatment, previous exposure to *Bd* and *Rv* concentration were significant (χ^2^_2_ = 7.85, *P* = 0.02; Fig. 2A). To scrutinise the pattern behind this interaction we separately analysed the treatment groups exposed to the low and the high *Rv* concentration. In the treatment groups exposed to the low *Rv* concentration, the interaction between previous exposure to *Bd* and thermal treatment was significant (χ^2^_2_ = 13.12, *P* = 0.001), where *Rv* infection intensity tended to be higher in the previously *Bd*-exposed treatment groups, but this effect was abolished by the 30°C thermal treatment (Fig. 2A; Table S5). In treatment groups exposed to the high *Rv* concentration, the two-way interaction between previous exposure to *Bd* and thermal treatment did not reach significance (χ^2^_2_ = 3.48, *P* = 0.18), and previous exposure to *Bd* did not have an effect (χ^2^_1_ = 1.49, *P* = 0.22), but thermal treatment did (χ^2^_2_ = 14.29, *P* < 0.001). Tadpoles in the 30°C treatment exhibited lower *Rv* copy numbers than those maintained at 22°C (Fig. 2A; Table S6).

Survival of agile frog tadpoles was significantly influenced by thermal treatments (COXME; χ^2^_2_ = 11.48, *P* = 0.003): it increased to 61.5% at 30°C from ca. 43.5% at 22-28°C (Fig. S2; Table S4). The effect of *Rv* concentration was also significant (χ^2^_1_ = 52.83, *P* < 0.001): survival probability was 2.7 times higher in the *Rv*-low treatment than in the *Rv*-high treatment (Fig. S2; Table S4). Previous co-exposure to *Bd* did not affect survival (χ^2^_1_ = 1.24, *P* = 0.27) and none of the interactions were significant (all *P* > 0.27).

### Common toads

The initial prevalence of *Bd* varied between 5 and 42%, and *Bd* infection intensities remained low until the start of thermal treatments (Table S2). At the same time, the initial prevalence of *Rv* varied between 20 and 100% and initial *Rv* infection intensities were low to moderate (Table S2; for details, see the Supplementary material).

After thermal treatments, the prevalence of *Bd* in *Bd*-exposed common toad tadpoles varied between 0 and 45% (Table S3). The thermal treatment (GLMM; χ^2^_2_ = 10.943, *P* = 0.004) significantly influenced *Bd* prevalence, but the presence and concentration of *Rv* did not (χ^2^_2_ = 2.379, *P* = 0.304; Fig. 2C). The prevalence of *Bd* was higher in tadpoles treated at 22°C compared to those kept at 30°C (Table S7). The interaction between thermal treatment and the presence and concentration of *Rv* was non-significant (χ^2^_4_ = 3.681, *P* = 0.45).

The intensity of infection with *Bd* was highest after 22 °C treatments in all *Bd*-exposed groups, with a dramatic drop in GE values in groups receiving 28 and 30°C treatments (Table S3; Fig. 2C). The effect of co-exposure to *Rv* could not be assessed across all thermal treatments due to the very low prevalence observed at higher temperatures, but within the 22 °C thermal treatment it was non-significant (χ^2^_2_ = 0.065, *P* = 0.968).

The prevalence of *Rv* in *Rv*-exposed tadpoles varied between 0 and 40% (Table S3, Fig. 2B) and was significantly influenced by thermal treatment (χ^2^_2_ = 11.92, *P* = 0.002), *Bd* co-exposure (χ^2^_1_ = 5.43, *P* = 0.02) and *Rv* concentration (χ^2^_1_ = 8.84, *P* = 0.003). The prevalence of *Rv* was higher when tadpoles were treated at 22°C compared to 28 or 30°C, while it did not differ between animals treated at 28 and 30°C (Table S7; Fig. 2B). Also, *Rv* prevalence was higher in the absence of *Bd* co-exposure and after receiving a high *Rv* concentration (Table S7; Fig. S3). Copy number of *Rv* was highest after the 22°C treatment, especially when tadpoles had been exposed to the high *Rv* concentration in the absence of *Bd* (Fig. 2B).

## Discussion

Previous reports showed that interactions between *Bd* and *Rv* during co-infection can be negative and positive and also can result in neutral co-existence [11]. Under natural circumstances, the secondary invader may encounter already triggered immune functions of the host, resulting in lowered replication. On the other hand, if the pathogen arriving second faces immune responses that are weakened by infection by the preceding pathogen, replication of the secondary agent may be facilitated [26]. In the present study, previous exposure to *Bd* decreased *Rv* prevalence after a week in both species. In contrast, *Bd* prevalence was not consistently affected by a single exposure to *Rv* that followed several weeks of *Bd* exposure. However, some effects of co-infection were modulated by the concentration of *Rv* inoculation and the subsequent thermal treatment. Generally, high *Rv* concentrations resulted in enhanced *Rv* prevalence in both hosts and higher viral loads in agile frogs. The dose of inoculum is a crucial factor that increases virulence and decreases average survival time towards high doses in amphibians [56], which corresponds to our findings. The other general trend in our results was that the replication of both pathogens was lowest at 30°C, resulting in the lowest prevalence and infection intensities. The outcome of these three effects (i.e., co-infection, *Rv* concentration, and temperature) was complex. In agile frogs, *Bd* co-exposure increased *Rv* replication in the low-*Rv* treatment at lower temperatures, but this effect was reversed at 30°C. This pattern was not observed in agile frog tadpoles exposed to high *Rv* concentration. In common toads, both high-temperature treatments resulted in zero *Rv* prevalence in all groups excepting the combination of high *Rv* concentration and absence of *Bd* exposure. These findings highlight that co-infections can alter disease dynamics and they may do so in a temperature-dependent way.

Our results indicate that the thermal tolerance of the two pathogens might not be as different as previously thought. In the case of single *Bd* infections, a series of *in vivo* studies performed on larval and adult frogs demonstrated that elevated temperatures (approx. 26-30°C) could reduce *Bd* growth, enhance survival or clear the pathogen burden depending on the host species, the applied temperature and the duration of thermal treatment [41-43]. In contrast, the thermophilic nature of *Rv* documented by *in vitro* studies [33] and supported by the observation that mortality is highest in the warm summer months [34] put forward the hypothesis that *Rv* would become more virulent at higher temperatures, irrespective of co-infection with *Bd*. In contrast to this prediction, in our study, both host species exhibited lowered infection prevalence and intensity of *Rv* at 30°C. This had a crucial effect on fitness because, among the agile frog tadpoles exposed to *Rv*, those treated at 30°C had the lowest mortality. These findings have several implications for important conservation issues. First, temperature variability associated with anthropogenic climate change is one of the most current problems that can dramatically impact wildlife. More specifically, the interactions between climate warming and disease outbreaks have caused declines or even extinctions in several ectothermic hosts, including amphibians [57-59]. Our study found direct experimental evidence that if the temperature becomes higher than what is ideal for the pathogens (i.e., close to 30°C), it can reduce disease risk in amphibian larvae under simultaneous threat to two widespread pathogens. Thus, our empirical findings of disease outcomes during co-infection with *Bd* and *Rv* might facilitate the parameterisation of predictive models aiming to forecast the consequences of changing climate for future dynamics of wildlife diseases [35, 60]. Incorporating co-infections into predictive models delivers higher predictive power than single-parasite models by improving estimates of co-infection prevalence at the individual level [61]. Finally, the artificial elevation of environmental temperature beyond the pathogens’ optimum could serve as a basis for *in situ* conservation actions against chytridiomycosis and ranavirosis [49].

Our study also revealed inter-specific differences in pathogen resistance and tolerance. We found that agile frogs were highly resistant to the chytrid fungus but susceptible to ranaviral infection. At the same time, common toads were moderately resistant to both pathogens. These differences in prevalence were mirrored by patterns in mortality. Agile frogs exposed to *Rv* suffered considerable and concentration-dependent mortality in accordance with other amphibian-ranavirus systems [62]: tadpoles that received high concentrations were less likely to survive than tadpoles challenged with low concentrations [56]. In contrast, common toads experienced negligible mortality regardless of pathogen exposure. A previous experiment investigating the chemical defences against *Bd* reported a similarly low susceptibility to the fungus in the early life stages of agile frogs compared to common toads [63]. The susceptibility of amphibian species to chytridiomycosis has been related to the presence/absence of cytolytic skin-secreted antimicrobial peptide (AMP) profiles [64, 65]. Accordingly, the resistance of agile frogs to *Bd* might be related to their AMP production (e.g., Brevinin-1 Da [66]). However, as agile frogs in our study carried *Rv* at high prevalence, this line of defence appeared ineffective against *Rv* infection. Antimicrobial peptides can directly inactivate plaque formation of FV3 *in vitro*, but they do not inhibit viral replication in infected cells [67]. In contrast, bufonid toads lack skin-secreted AMPs [68], but from early larval development, they produce bufadienolide compounds [69] with antimicrobial activity, which can protect against *Bd* [70, 71] and might protect against *Rv*. Furthermore, while AMPs are present exclusively on skin surfaces and may be effective against skin invaders such as the chytrid fungus, bufadienolides are present not only in the skin but also in internal organs [72] and thus might more successfully mitigate pathogens that target internal organs such as *Rv*. The effects of bufadienolides on *Rv* replication or co-infection with other pathogens are not yet explored but may hold promising potential for battling diseases.

In summary, our results suggest that high temperatures may be beneficial to amphibians exposed to both *Bd* and *Rv*. Also, while previous exposure to *Bd* affected *Rv* prevalence and in some treatment combinations *Rv* replication as well, superinfection with *Rv* did not influence the replication of *Bd*. Finally, temperature and co-infection appeared to also interact in their effects on pathogen replication and disease progression. Nonetheless, in our study, both species exhibited relatively low prevalence and infection intensities except for *Rv* in agile frogs, so we urge further experimental studies on more susceptible species to scrutinise the effects of co-infection and external factors modulating its outcomes in amphibians.

## Summary

Rising temperatures can facilitate epizootic outbreaks, but disease outbreaks may be suppressed if temperatures increase beyond the optimum of the pathogens while still within the temperature range that allows for effective immune function in hosts. The two most devastating pathogens of wild amphibians, *Batrachochytrium dendrobatidis* (*Bd*) and ranaviruses (*Rv*), co-occur in large areas, yet little is known about the consequences of their co-infection and how these consequences depend on temperature. Here we tested how co-infection and elevated temperatures (28 and 30°C vs. 22°C) affected *Bd* and *Rv* prevalence, infection intensities, and resulting mortalities in larval agile frogs and common toads. We found multiple pieces of evidence that the presence of one pathogen influenced the prevalence and/or infection intensity of the other pathogen in both species, depending on temperature and initial *Rv* concentration. Generally, the 30°C treatment lowered the prevalence and infection intensity of both pathogens, and, in agile frogs, this was mirrored by higher survival. These results suggest that if temperatures naturally increase or are artificially elevated beyond what is ideal for both *Bd* and *Rv*, amphibians may be able to control infections and survive even the simultaneous presence of their most dangerous pathogenic enemies.

## Acknowledgements

We thank M. Szederkényi for field and technical assistance and for Z. Boros, C. Kalina, R. Bertalan, B. Üveges, N. Ujhegyi and A. Merkei for their help in conducting experiments. The authors are indebted to V. Krízsik and M. Tuschek (Molecular Taxonomy Laboratory, NHMUS), K. Ursu (NÉBIH) and M. Németh (ATK NÖVI) for their technical assistance during laboratory work. We thank A. Doszpoly (ÁOTKI, ELKH) for help with *Ranavirus* maintenance. The *Bd*-GPL lineage used in this study was obtained from T.W.J. Garner (ZSL) and zoospore genomic equivalents kindly provided by J. Bosch (MNCN). The Frog Virus 3 isolate was obtained from R.E. Marschang (Laboklin GmbH). We thank B. Bombay for the tadpole paintings.

## Declarations

### Funding

The study was funded by the National Research, Development and Innovation Office of Hungary (NKFIH, grant K-124375 for AH and grant NN-140356 for TP). The authors were supported by the New National Excellence Program of the Ministry for Innovation and Technology of the National Research, Development and Innovation Fund (ÚNKP-19-3, ÚNKP-20-3 and ÚNKP-21-3 to AK, ÚNKP-19-4 to AH and ÚNKP-21-4 to UJ). DHerczeg was supported by the Young Investigators Programme of the Hungarian Academy of Sciences and VB by the János Bolyai Research Scholarship.

### Conflict of interest

The authors declare no conflict of interest.

### Ethical approval

All experimental procedures were approved by the Ethical Commission of the ELKH ATK NÖVI under Good Scientific Practice guidelines and national legislation. All experiments were carried out according to the permits issued by the Government Agency of Pest County (Department of Environmental Protection and Nature Conservation, PE/EA/295-31 7/2018 and PE/EA/58-4/2019).

### Authors’ Contributions

DHerczeg and AH conceived and designed the experiments. DHerczeg, DHolly, AK and JU performed the experiments. DHerczeg, DHolly, AK, TP, HT-V, and JU performed the molecular analysis. DHerczeg and VB analysed the data. DHerczeg, VB and AH wrote the manuscript; other authors provided editorial advice.

## Electronic supplementary material

### Supplement to the description of methods

#### Animal collection and husbandry

In March and April 2019, we collected ca. 100 eggs from each of ten freshly laid clutches of agile frogs and common toads from natural populations located in and around Budapest, Hungary. We sampled five agile frog egg clutches collected from Ilona-tó: 47.71326 N, 19.04050 E and five from Hortobai-katlan: 47.71110 N, 19.04570 E, while five common toad egg strings from Apát-kúti-tározó: 47.774736 N, 18.986300 E and five from Hidegkúti horgásztó: 47.569131 N, 18.954631 E. We transported the eggs to the Júliannamajor Experimental Station of the Plant Protection Institute, Centre for Agricultural Research in Budapest (ATK NÖVI; 47.547778 N, 18.934722 E).

We reared the embryos separated according to sib-groups (families) in plastic boxes (25 × 17 × 13 cm) holding 1-L reconstituted soft water (RSW) and maintained a temperature of 16.3 ± 0.3°C (mean ± SD) and a 12:12 h light:dark cycle. Five days after hatching, the laboratory air temperature was raised to 18.45 ± 0.5°C (mean ± SD), and we maintained a 13:11 h light:dark cycle. At this time, each sib-group was haphazardly divided into groups of 10 larvae and placed into plastic rearing containers (37 × 27 × 16.5 cm), each holding 10 litres of reconstituted soft water (RSW). We changed the RSW twice a week and fed tadpoles with chopped and slightly boiled spinach *ad libitum*. Infection with *Batrachochytrium dendrobatidis* (*Bd*) was applied at each of the first five water changes as described below (“*Experimental infections*”).

Four days after the 5^th^ *Bd* inoculation, on day 19, we haphazardly selected eight tadpoles from each rearing container, placed them individually into 2-L plastic rearing boxes filled with 1.8 litres of RSW and randomly assigned them to *Ranavirus* (*Rv*) × temperature treatment combinations. We placed rearing boxes on a shelf system in randomised spatial blocks containing one replicate from each treatment combination. The tadpoles were housed in these individual boxes for a short time before the 24-hours *Rv* treatment (as described below; see “*Experimental infections*”) and during the 6-days thermal treatment (see “*Thermal treatments*” below).

#### Experimental infections

We used a liquid culture of the *Bd* isolate IA042 (obtained from a dead *Alytes obstetricans* in 2004 in Spain), which belongs to the global pandemic lineage (*Bd*-GPL). We maintained the stock culture in mTGhL broth (8 g tryptone, 2 g gelatine hydrolysate, and 4 g lactose in 1000 ml distilled water) in 25 cm^2^ cell culture flasks at 4°C and passaged every three months. One week before use in the experiment, we inoculated 100 ml mTGhL with 2 ml stock culture in 175 cm^2^ flasks and incubated these cultures at 20°C for seven days. We estimated *Bd*-zoospore concentrations using a Bürker chamber at ×400 magnification and inoculated each rearing container assigned to the *Bd* and the *Bd* + *Rv* treatments with 10 ml to obtain final concentrations of approximately 2000 zoospores×ml^-1^.

The FV3 strain was propagated in T75, and T175 cell-culture flasks on Epithelioma Papulosum Cyprini (EPC) cells (ATCC CRL-2872) in DMEM nutrient medium supplemented with 2% fetal bovine serum (FBS), without the use of antibiotics at 25°C with 5% CO_2_. Flasks were freeze-thaw harvested, and the medium was clarified from cell-debris by centrifugation for 10 min at 2500 × g to collect a sufficient amount of viral supernatant. Viral titre (TCID_50_) was determined on 96-well plates of EPC cells using the Spaerman-Karber formula and converted to plaque-forming units (pfu) using the equation: 1 TCID_50_ = 0.69 pfu. We exposed tadpoles to FV3 by applying one of two concentrations (i.e., *Rv*-low and *Rv*-high) during 24-hour exposures immediately preceding the start of the thermal treatment. We placed tadpoles allocated to *Rv* treatments individually in 500 mL plastic cups containing 200 mL RSW and 6.12 × 10^3^ pfu×ml^-1^ FV3 in the *Rv*-low group and 6.25 × 10^5^ pfu×ml^-1^ FV3 in the *Rv*-high group. We added the same quantity of sham extract (only the DMEM nutrient medium with 2% FBS without the virus) to the rest of the tadpoles.

#### Thermal treatments

We placed the 2-L rearing boxes in 80 × 60 × 12 cm trays filled with tap water to a depth of 8 cm (water level was lower than in rearing boxes by ca. 2 cm to avoid floating of the latter) and subsequently turning on submersible aquarium heaters (Tetra HT 200 in 28°C treatments and Tetra HT 300 in 30°C treatments) and water pumps (Tetra WP 300) placed opposite to each other on the longitudinal axis of trays. Thereby, water temperature increased gradually to the desired level in ca. two hours, allowing tadpoles to adjust to increasing temperatures. After heating up, the temperature did not change over time and varied only a little among/within trays (Table S1), as documented by automated temperature loggers (Onset HOBO Pendant Temperature/Light 8K) placed into one-third of the trays (i.e., 12 out of 36). Actual water temperatures in the tadpole rearing boxes were overall 21.4 ± 0.72, 28.16 ± 0.24 and 30.13 ± 0.35°C (mean ± SD) in the three temperature treatments, respectively.

During the six days of thermal treatment, we changed water twice with RSW pre-heated to the temperature of the respective thermal treatment group. We fed tadpoles with a lowered amount of spinach (one-third of the amount provided during the rearing period) to avoid water fouling and anoxia at high temperatures.

#### Molecular analyses

We assessed *Bd* infection intensity from dissected mouthparts in the case of *Bd* and from liver tissue in the case of *Rv*. The small body size of common toad tadpoles did not allow for precise separation of the liver, so in their case, we used all internal organs as a whole. We homogenised mouthparts with Qiagen Tissue Lyser II (Qiagen, Hilden, Germany) and the liver or all internal organs with a disposable pellet mixer (VWR, catalog no. 47747-370). The *Bd* DNA was extracted with PrepMan Ultra Sample Preparation Reagent (Thermo Fisher Scientific, Waltham, Massachusetts, USA), and the *Rv* DNA with Wizard Genomic DNA Purification Kit (Promega, Madison, Wisconsin, USA) according to the manufacturer’s protocol. We stored extracted DNA at −20°C until further analyses.

We assessed *Bd* infection prevalence and intensity using qPCR following a standard amplification methodology targeting the ITS-1/5.8S rDNA region. To avoid PCR inhibition by ingredients of PrepMan, we diluted samples ten-fold with double-distilled water. We ran samples in duplicate, and when the result was equivocal, we repeated reactions in duplicate. If we obtained an equivocal result again, we considered the sample *Bd* positive. Genomic equivalent (GE) values were estimated from standard curves based on four standard dilutions (100, 10, 1 and 0.1 zoospore genomic equivalents). We estimated *Rv* infection prevalence and intensity by targeting the major capsid protein (MCP) gene of the viral genome. We ran samples in duplicate and followed the protocol detailed above in case of equivocal results. All reactions were run on a BioRad CFX96 Touch Real-Time PCR System.

#### Statistical analyses

We calculated the prevalence data with 95% confidence intervals presented in Table S2 and Table S3 using QPWeb version 1.0.15 [73]. For all other analyses, we used the R computing environment, version 4.0.4 [74].

Before thermal treatment, *Bd* prevalence was low in both species, so we did not apply statistical analyses to initial *Bd* prevalence and infection intensity. For initial *Rv* prevalence, we tested the effects of the applied *Rv* concentration and the previous exposure to *Bd* by Fisher’s Exact Tests. For initial *Rv* infection intensity, we used linear mixed-effects models (LMM; ‘lme’ function of the ‘nlme’ package) to simultaneously analyse the effects of *Rv* concentration and *Bd* co-infection and their interaction.

To analyse prevalence and pathogen load after thermal treatment, we included only those treatment groups that had been exposed to the given pathogen. We used generalised linear mixed-effects models (GLMM) to test the effects of treatments on pathogen prevalence. The model for *Rv* prevalence in agile frogs contained thermal treatment (22, 28 or 30°C), *Rv* concentration (low or high), co-exposure with *Bd* (yes or no) as fixed factors and all two-way interactions (there was not enough variance in the data to allow model fit for testing the three-way interaction). The model for *Bd* prevalence in common toads contained thermal treatment and a three-category factor that combined the information on the presence and concentration of *Rv* (no *Rv*, low *Rv*, high *Rv*) and the interaction of the two fixed factors. In both models, we entered family as a random factor. We assumed a binomial error distribution and used a logit link function. We fitted the models applying maximum likelihood estimation using the ‘glmmTMB’ function of the package ‘glmmTMB’ [75] and checked model-fit diagnostics using the ‘DHARMa’ package [76]. We did not run such analyses for *Bd* prevalence in agile frogs because zero prevalence in the majority of treatment groups would have led to very high estimation uncertainty (separation) and inability to test interactions.

We analysed the effect of treatments on infection intensity after temperature treatment within *Rv*-positive agile frogs using a linear mixed-effects model, allowing the variances to differ among treatment groups (‘varIdent’ function) because graphical model diagnostics indicated heterogeneous variances. The model contained the natural log-transformed *Rv* infection intensity as a dependent variable, thermal treatment, *Rv* concentration, *Bd* co-exposure and their two-and three-way interactions as fixed factors, and family as a random factor. For *Rv* infection load in common toads, we used the same modelling approach, but we tested only the main effects because there was not enough variation in the data for testing interactions (i.e. prevalence was zero in 6 out of 12 treatment combinations, causing separation in binomial models). Low prevalence prohibited the analyses of infection intensity for *Bd* in both species.

To analyse the survival of agile frog tadpoles during thermal treatment, we ran a mixed-effects Cox’s proportional hazards model (COXME; ‘coxme’ function of the ‘coxme’ package), entering family as a random effect [77]. We entered survival as an ordinal categorical dependent variable ranging 1-6 (each category representing the day of death during heat treatment, 1 being the first 24 hours); individuals that survived to the end of thermal treatment were treated as censored observations. We included thermal treatment, *Rv* concentration, *Bd* co-exposure and their two- and three-way interactions as predictors. Because mortality of common toads was negligible (1.1%; see Table S3), we did not analyse their survival.

We applied a backward stepwise model selection procedure to reduce noise in parameter estimates due to the inclusion of non-significant terms [78, 79]. We obtained statistics for excluded terms by re-entering them to the final model. For these steps, we used type-3 analysis-of-deviance tables (‘Anova’ function of the ‘car’ package). To perform pairwise comparisons, we calculated linear contrasts from the final models using the ‘emmeans’ function of the ‘emmeans’ package while applying the false discovery rate (FDR) correction method to adjust *P* values for multiple comparisons [80, 81].

## Supplementary results

### Prevalence and infection intensity of pathogens before the thermal treatment

Cross-contamination was not detected in either infection group except for one individual in the control group after 30°C thermal treatment, but this tadpole also exhibited very low *Rv* infection intensity (28 pfu×ml^-1^). Seven agile frog samples collected after thermal treatments were accidentally lost during sample procession, so that the sample sizes in the infection groups were N_control_ = 59; N_*Bd*_ = 58; N_*Rv*-low_ = 58; N_*Rv*-high_ = 59; N_*Bd*+*Rv*-low_ = 60; N_*Bd*+*Rv*-high_ = 59; resulting in a sample size of 353 in agile frogs. We lost one common toad sample after the thermal treatment from the uninfected control group (N_control_ = 59), resulting in a total sample size of 359 in common toads. Four samples preserved after experimental infections but before thermal treatments exhibited low DNA quality (one in the low concentration *Rv* treatment and one in the high concentration *Rv* treatment in the case of agile frogs; and one in the low concentration *Rv* treatment, and one in the *Bd* + high concentration *Rv* treatment in case of common toads), so we excluded these from the analysis of initial infection, resulting in a sample size of 118 individuals in both species.

All but two of the 120 agile frog tadpoles survived until the start of thermal treatment. At that time, the initial prevalence of *Bd* in *Bd*-exposed individuals was extremely low: two out of 60 (Table S2). Initial *Bd* infection intensities were also very low, the Genomic Equivalent (GE) values being 0.36 and 2.43, respectively (Table S2). In contrast, the initial prevalence of *Rv* in *Rv*-exposed tadpoles was high, varying between 73.7 and 100% (Table S2), and was significantly higher after exposure to high *Rv* concentration compared to low *Rv* concentration (Fisher’s Exact Test; *P* = 0.005), while the presence of *Bd* had no effect on *Rv* prevalence (Fisher’s Exact Test; *P* = 0.45). Initial *Rv* infection intensities within the infected agile frog tadpoles were low to moderate (Table S2) and significantly higher in the *Rv*-high group than in the *Rv*-low group (LMM; χ^2^_1_ = 127.612, *P* < 0.001), while the presence of *Bd* had no effect (χ^2^_1_ = 0.003, *P* = 0.951). The interaction between the presence of *Bd* and the concentration of *Rv* was non-significant (χ^2^_1_ = 0.03, *P* = 0.861).

All but two of the 120 common toad tadpoles survived until the thermal treatment. The initial prevalence of *Bd* in *Bd*-exposed individuals varied between 5 and 42% (Table S2). The initial prevalence of *Rv* in *Rv*-exposed tadpoles varied between 20 and 100% (Table S2), and it was significantly higher after exposure to high *Rv* concentration compared to low *Rv* concentration (Fisher’s Exact Test; *P* < 0.005), while the presence of *Bd* had no effect on *Rv* prevalence (Fisher’s Exact Test; *P* = 0.45). Also, the initial infection intensity of *Rv* in *Rv* exposed toad tadpoles was higher in individuals that received high *Rv* concentrations (LMM; χ^2^_1_ = 130.721, *P* < 0.001) without an effect of the presence of *Bd* (χ^2^ = 1.505, *P* = 0.219). The interaction between the presence of *Bd* and the concentration of *Rv* was non-significant (χ^2^ = 0.643, *P* = 0.422).

**Table S1.** Temperature data as documented by automated temperature loggers, i.e., four loggers assigned to each nominal temperature in heating trays during the thermal treatments. Note that the temperatures in heating trays were somewhat higher than the actual temperature in the tadpole rearing boxes during the thermal treatment.

| **Nominal**<br>**temperature**<br>**(°C)** | **Mean** | **SD** | **Range** | **SE** |
| --- | --- | --- | --- | --- |
| 22 | 21.83 | 0.36 | 1.53 | 0.02 |
| 22 | 22.39 | 0.4 | 1.63 | 0.02 |
| 22 | 22.62 | 0.32 | 1.34 | 0.01 |
| 22 | 22.43 | 0.47 | 1.82 | 0.02 |
| 28 | 28.83 | 0.37 | 1.7 | 0.02 |
| 28 | 29.30 | 0.21 | 1.2 | 0.01 |
| 28 | 28.81 | 0.36 | 1.49 | 0.02 |
| 28 | 29.72 | 0.45 | 1.5 | 0.02 |
| 30 | 31.65 | 0.37 | 1.74 | 0.02 |
| 30 | 31.23 | 0.27 | 1.02 | 0.01 |
| 30 | 31.03 | 0.27 | 1.02 | 0.01 |
| 30 | 31.53 | 0.37 | 3.14 | 0.02 |

**Table S2.** Initial prevalence (Prev) and infection intensity of *Batrachochytrium dendrobatidis* (*Bd*) and *Ranavirus* (*Rv*) by host species and infection group as assessed immediately before the thermal treatments. We calculated mean infection intensities (Mean inf int) and corresponding standard errors (SE) by including only qPCR-positive specimens. N_surv_: Number of tadpoles that survived the infection trials; 95% CI = 95% confidence intervals. There were no cross-contaminations.

|  | N<sub>start</sub> | N<sub>surv</sub> | *Bd* |  |  |  |  |  | *Rv* |  |  |  |  |  |
| --- | --- | --- | --- | --- | --- | --- | --- | --- | --- | --- | --- | --- | --- | --- |
|  |  |  | N<sub>inf</sub> | Prev (%) | 95% CI | Mean inf int | Range | SE | N<sub>inf</sub> | Prev (%) | 95% CI | Mean inf int | Range | SE |
| Agile frog |  |  |  |  |  |  |  |  |  |  |  |  |  |  |
| Control | 20 | 20 | 0 | 0 |  |  |  |  | 0 | 0 |  |  |  |  |
| *Bd* | 20 | 20 | 1 | 5 | 0.2-24.4 | 0.36 |  |  | 0 | 0 |  |  |  |  |
| *Rv*-low | 20 | 19 | 0 | 0 |  |  |  |  | 14 | 73.7 | 50-89 | 781 | 4856 | 363 |
| *Rv*-high | 20 | 19 | 0 | 0 |  |  |  |  | 19 | 100 | 82.4-100 | 63000 | 377000 | 22000 |
| *Bd* + *Rv*-low | 20 | 20 | 0 | 0 |  |  |  |  | 17 | 85 | 62.7-95.7 | 701 | 3362 | 261 |
| *Bd* + *Rv*-high | 20 | 20 | 1 | 5 | 0.2-24.4 | 2.43 |  |  | 20 | 100 | 83.3-100 | 32500 | 96800 | 5964 |
| Common toad |  |  |  |  |  |  |  |  |  |  |  |  |  |  |
| Control | 20 | 20 | 0 | 0 |  |  |  |  | 0 | 0 |  |  |  |  |
| *Bd* | 20 | 20 | 5 | 25 | 10.4-47.4 | 0.63 | 2.3 | 0.44 | 0 | 0 |  |  |  |  |
| *Rv*-low | 20 | 19 | 0 | 0 |  |  |  |  | 5 | 26.3 | 11-50 | 29 | 89 | 18 |
| *Rv*-high | 20 | 20 | 0 | 0 |  |  |  |  | 20 | 100 | 83.3-100 | 2866 | 8148 | 618 |
| *Bd* + *Rv*-low | 20 | 20 | 1 | 5 | 0.2-24.4 | 0.15 |  |  | 4 | 20 | 7.1-42.3 | 22.5 | 34 | 8 |
| *Bd* + *Rv*-high | 20 | 19 | 8 | 42.1 | 22.1-65.5 | 9.66 | 40.5 | 4.96 | 19 | 100 | 82.4-100 | 7193 | 40100 | 2309 |

**Table S3.**
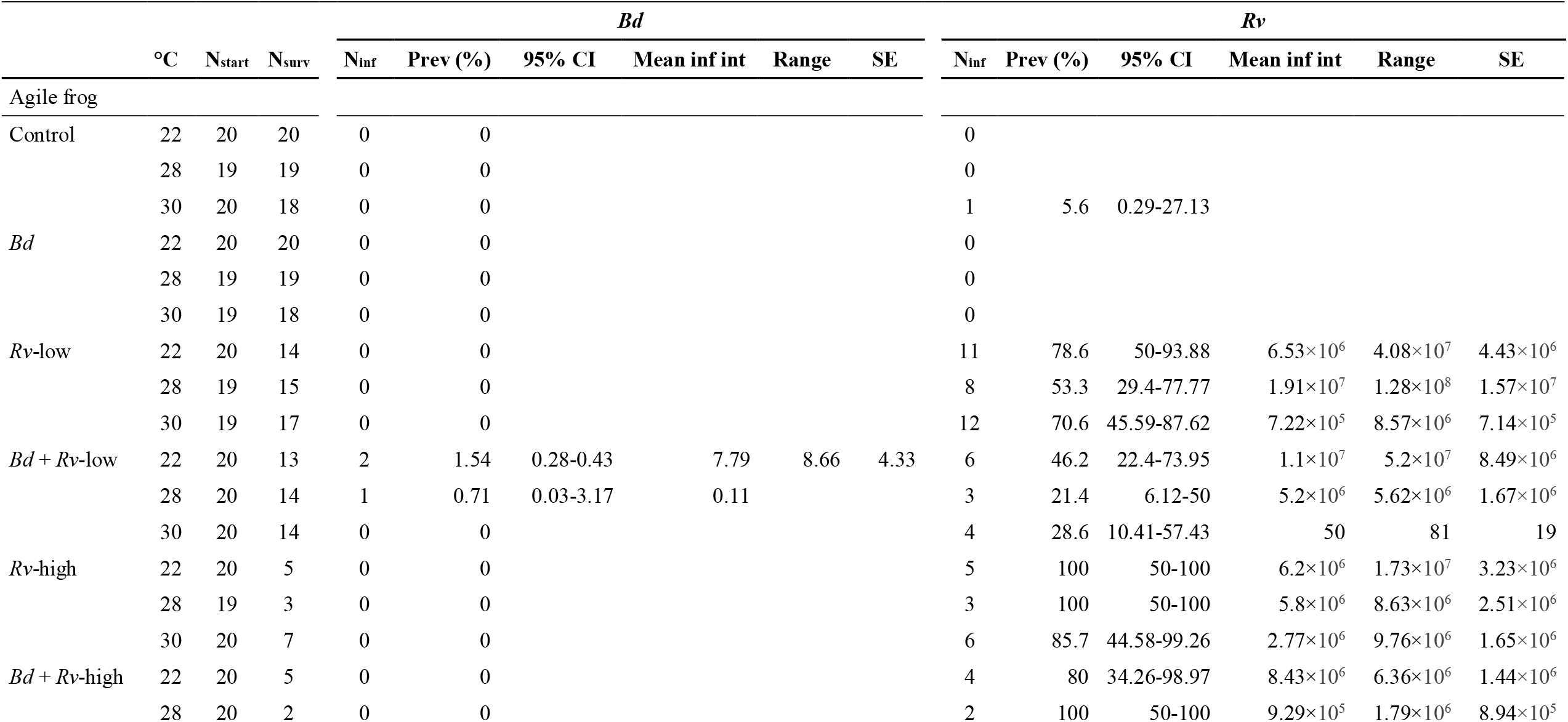
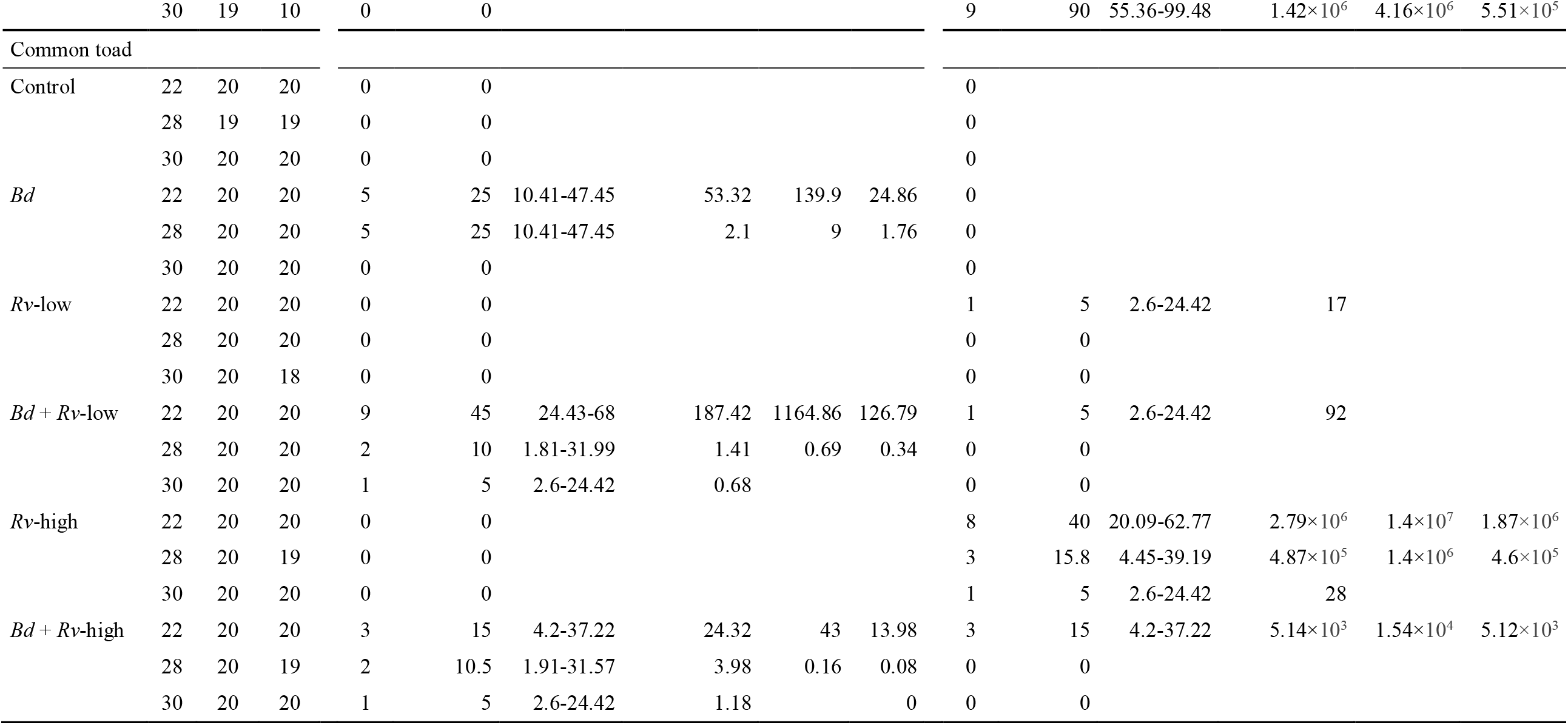
Prevalence (Prev) and infection intensity of *Batrachochytrium dendrobatidis* (*Bd*) and *Ranavirus* (*Rv*) by host species, infection group, and thermal treatment temperature (°C) as assessed immediately after the thermal treatments. We calculated mean infection intensities (Mean inf int) and corresponding standard errors (SE) by including only qPCR-positive specimens. N_start_: Number of tadpoles that entered the thermal treatments; N_surv_: Number of tadpoles surviving until sampling at the end of thermal treatment; 95% CI = 95% confidence intervals

**Table S4.** Agile frog responses to treatments. Linear contrasts (*c*), associated standard errors (SE), t-values (z-values in case of Cox’s proportional hazards model in the analysis of survival) and *P*-values adjusted using the FDR method are reported. Significant differences (*P* < 0.05) are in bold. Temp = Thermal treatment; *Bd* co-exp = Co-exposure to *Batrachochytrium dendrobatidis*; *Rv* conc = Concentration of *Ranavirus* exposure

| Dependent variable | Pairwise comparison | *c* | SE | *t* (or *z*) | *P* |
| --- | --- | --- | --- | --- | --- |
| *Rv* prevalence* | Temp 22 vs. 28 °C | 4.620 | 3.298 | 2.144 | 0.102 |
|  | Temp 22 vs. 30 °C | 2.268 | 1.466 | 1.267 | 0.282 |
|  | Temp 28 vs. 30 °C | 0.491 | 0.323 | -1.081 | 0.282 |
|  | ***Bd* co-exp no vs. yes** | **9.89** | **6.21** | **3.652** | **< 0.001** |
|  | ***Rv* conc high vs. low** | **27.2** | **23.7** | **3.807** | **< 0.001** |
| Survival** | Temp 22 vs. 28 °C | -0.281 | 0.219 | -1.284 | 0.199 |
|  | **Temp 22 vs. 30 °C** | **0.532** | **0.241** | **2.213** | **0.04** |
|  | **Temp 28 vs. 30 °C** | **0.813** | **0.241** | **3.376** | **0.002** |
|  | *Bd* co-exp no vs. yes | -0.208 | 0.187 | -1.113 | 0.265 |
|  | ***Rv* conc high vs. low** | **1.57** | **0.216** | **7.269** | **< 0.001** |

\*The linear contrast is the odds ratio
\*\*The linear contrast is the log (hazard ratio)

**Table S5.** Effect of *Bd* co-infection in each thermal treatment on *Ranavirus* (*Rv*) infection intensity of agile frogs exposed to low *Rv* concentration, and the difference of this effect between 30°C and the two lower temperatures. Linear contrasts (*c*, expressed as proportional difference: *Bd* present / *Bd* absent), associated standard errors (SE), t-values and *P*-values adjusted using the FDR method are reported. Significant differences (*P* < 0.05) are in bold. Temp = Thermal treatment; *Bd* co-exp = Co-exposure to *Batrachochytrium dendrobatidis*

| Temperature (°C) | *c* | SE | *t* | *P* |
| --- | --- | --- | --- | --- |
| 22 | 82.4 | 225.0 | 1.615 | 0.117 |
| **28** | **663.0** | **1661.4** | **2.593** | **0.044** |
| 30 | 0.1 | 0.1 | -1.951 | 0.091 |
| **22 & 28 vs. 30\*** | **2327** | **5110** | **3.530** | **0.001** |

\**P*-value is not adjusted because this is a contrast of contrasts

**Table S6.**
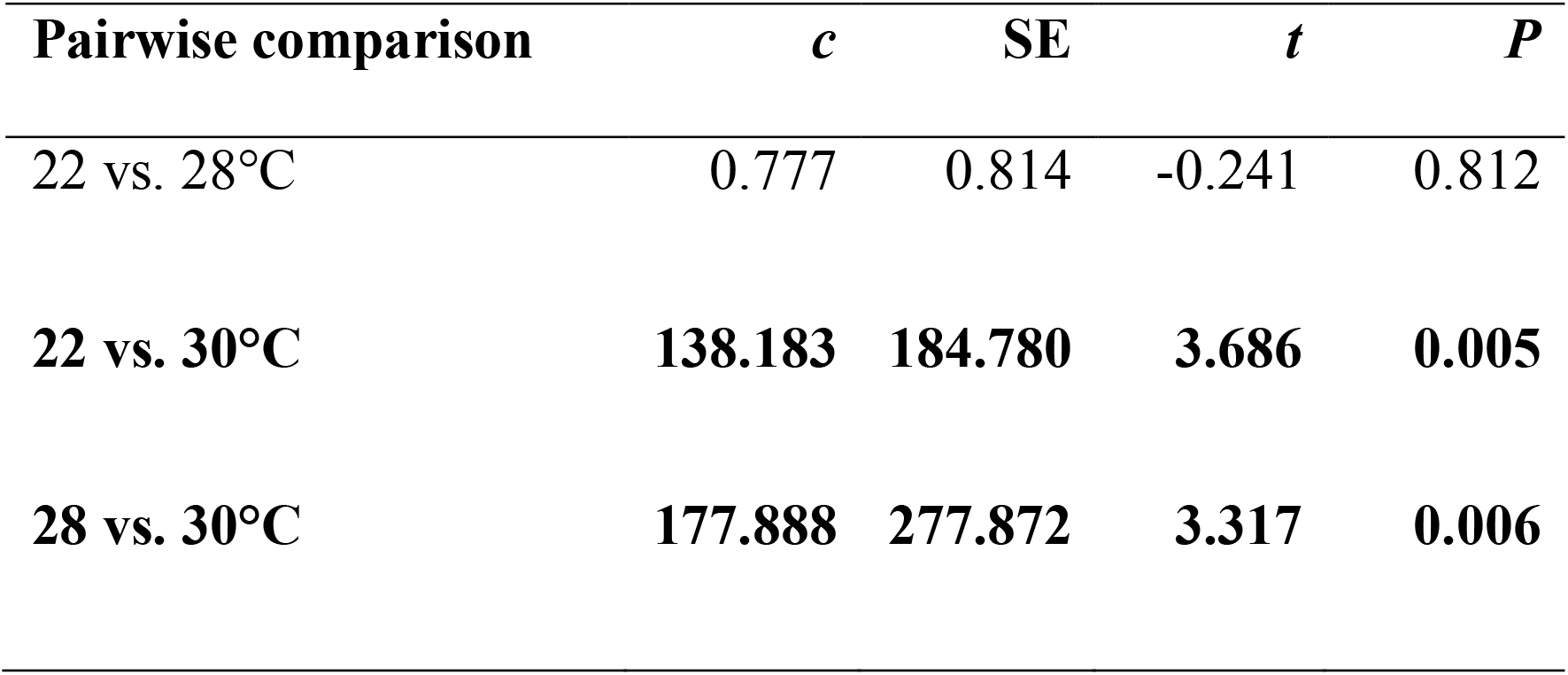
Differences between thermal treatment groups in *Ranavirus* infection intensity of agile frogs exposed to high *Rv* concentration. Linear contrasts (*c*, expressed as proportional difference between two thermal treatment groups), associated standard errors (SE), t-values and *P*-values adjusted using the FDR method are reported. Significant differences (*P* < 0.05) are in bold.

**Table S7.**
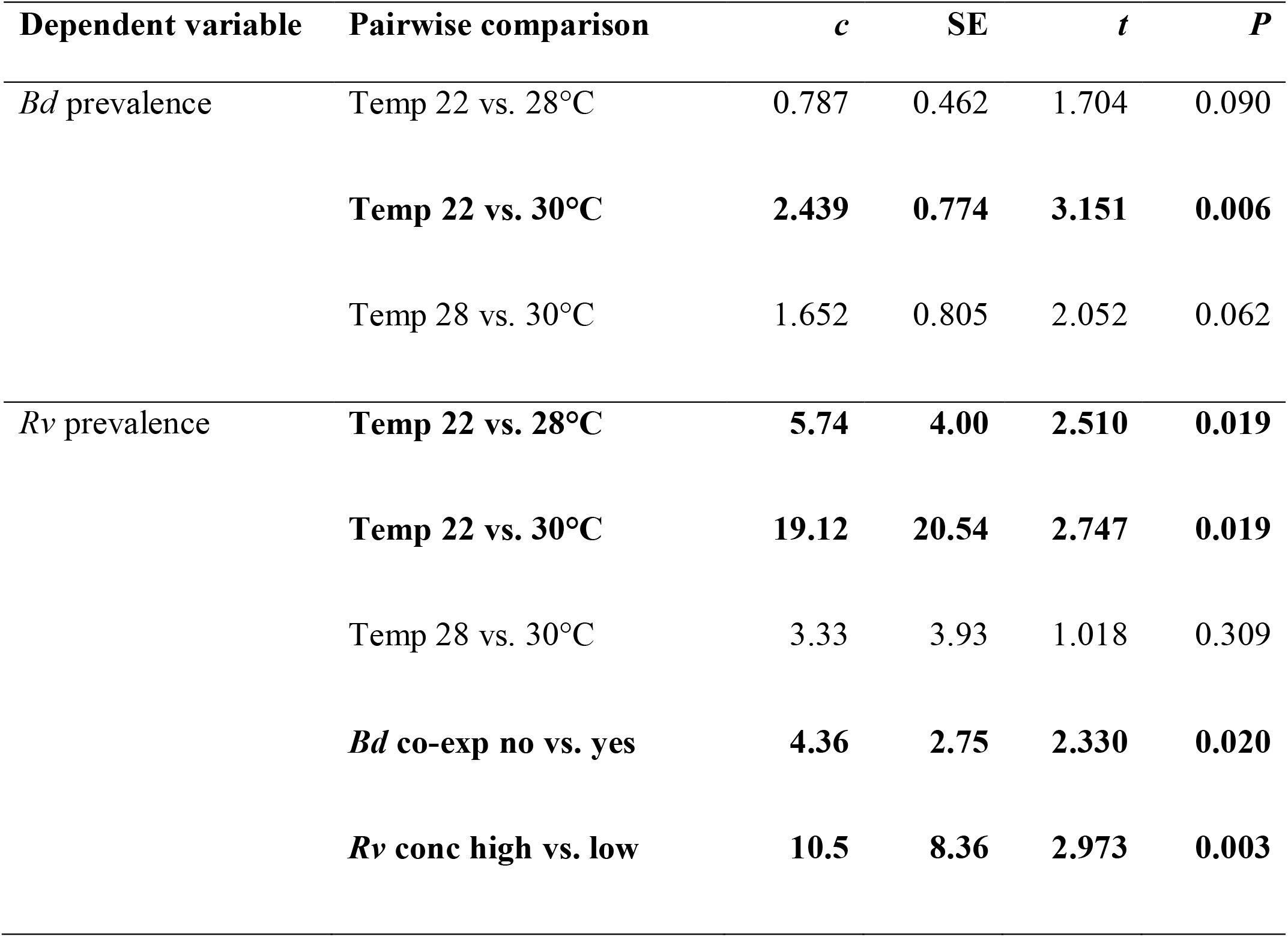
Common toad responses to treatments. Linear contrasts (*c*), associated standard errors (SE), t-values and *P*-values adjusted using the FDR method are reported. The linear contrast is the odds ratio. Significant differences (*P* < 0.05) are in bold. Temp = Thermal treatment; *Bd* co-exp = Co-exposure to *Batrachochytrium dendrobatidis*; *Rv* conc = Concentration of *Ranavirus* exposure

**Fig. S1.**
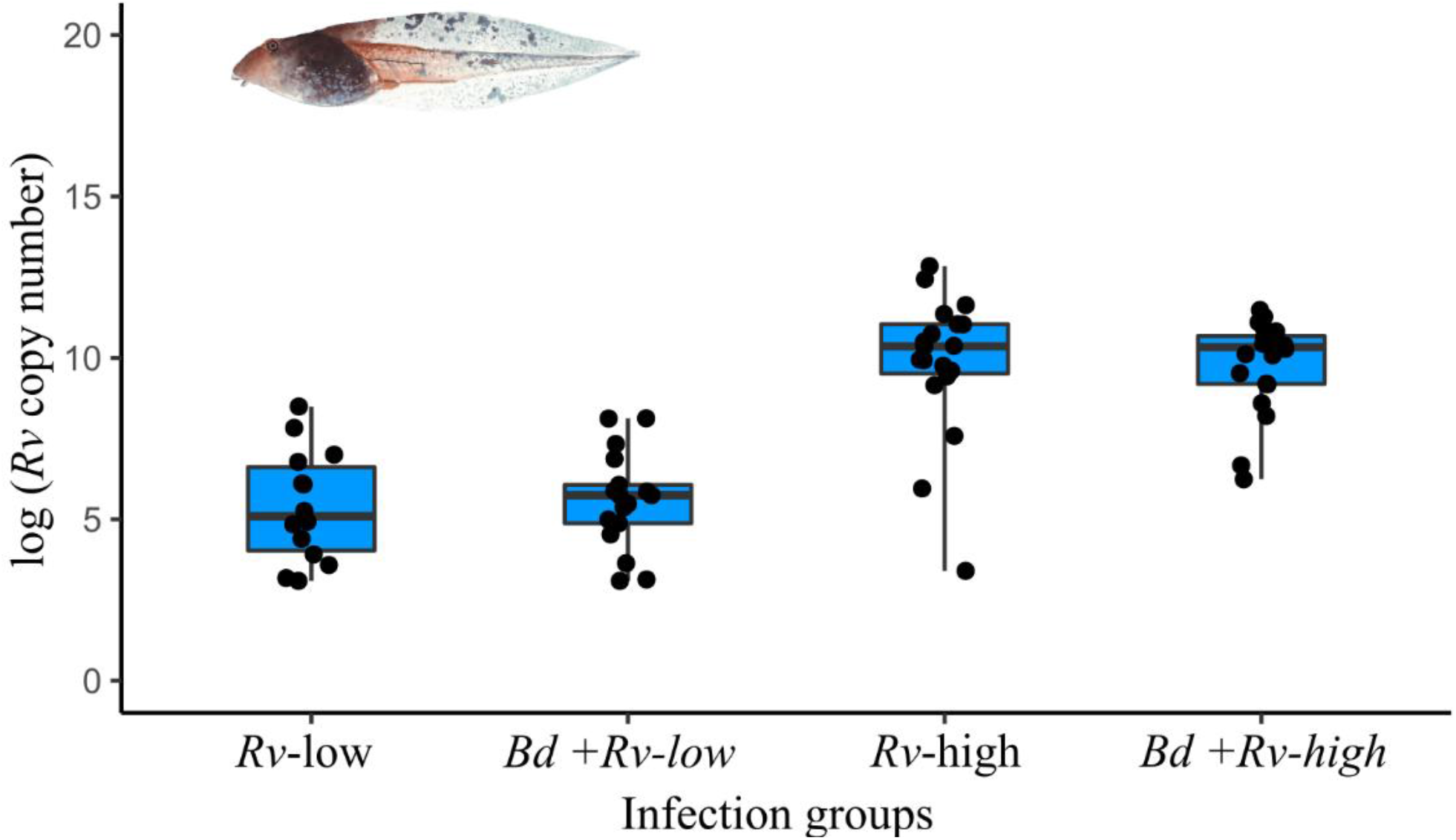
*Ranavirus* (*Rv*) infection intensities in *Rv*-positive agile frog tadpoles before the 6- days of thermal treatment, following exposure to low (*Rv*-low) or high (*Rv*-high) *Rv* concentration with or without co-exposure to *Bd* (*Batrachochytrium dendrobatidis*). The *Rv* infection intensity data (black dots) were natural log-transformed. Horizontal lines represent medians, boxes represent interquartile ranges, and whiskers represent minimum-maximum ranges.

**Fig. S2.**
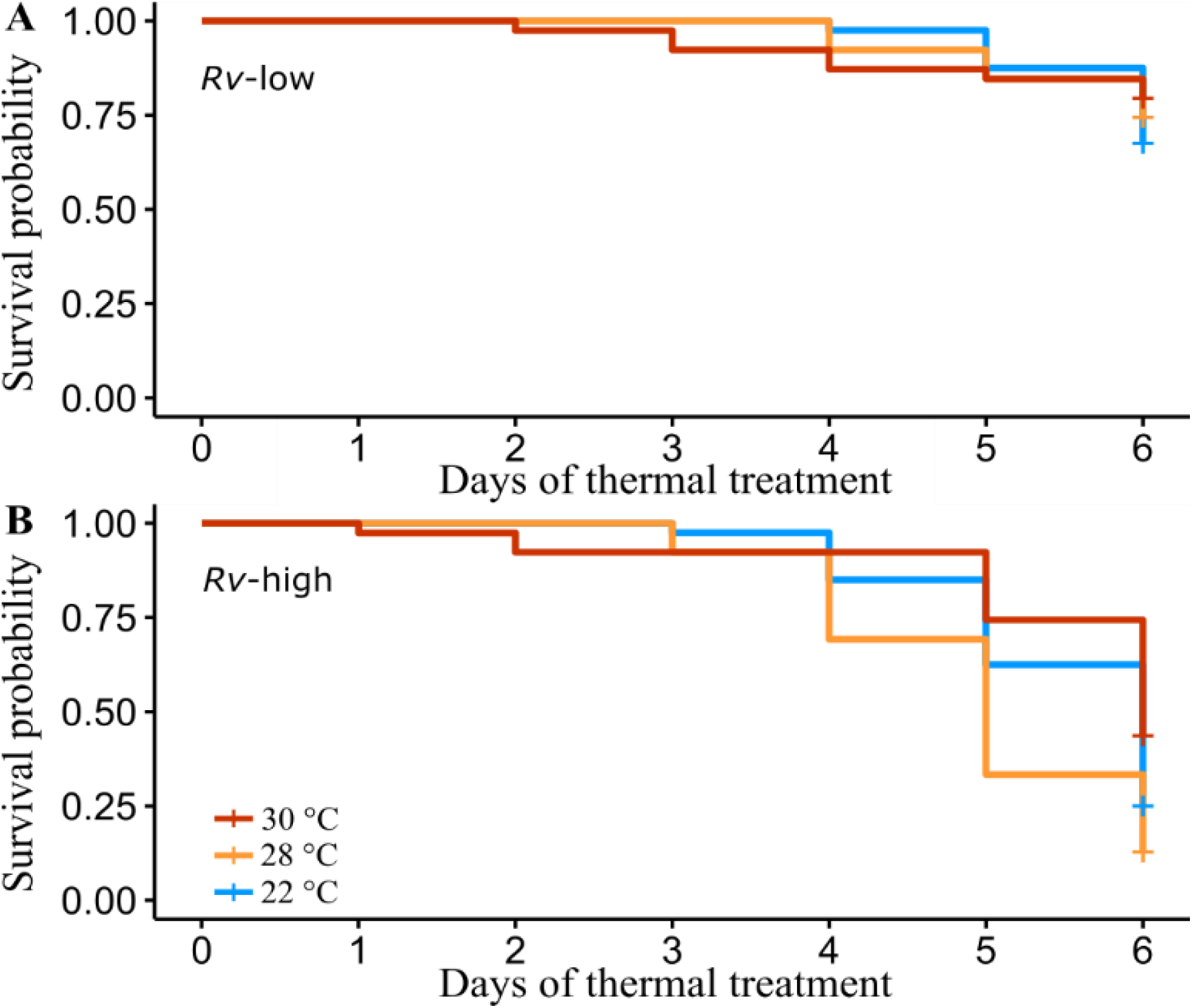
Survival probability of agile frog tadpoles (A) exposed to low *Rv* concentrations or (B) exposed to high *Rv* concentrations during the 6-days of thermal treatment visualised by Kaplan-Meier curves.

## Notes

### Competing Interest Statement

The authors have declared no competing interest.

